# High molecular weight hyaluronan – a potential adjuvant to fluid resuscitation in porcine abdominal sepsis

**DOI:** 10.1101/2021.08.12.456152

**Authors:** Annelie Barrueta Tenhunen, Jaap van der Heijden, Wojciech Weigl, Robert Frithiof, Paul Skorup, Anders Larsson, Anders Larsson, Jyrki Tenhunen

**Affiliations:** Hedenstierna Laboratory, Dept of Surgical Sciences, Uppsala University, Uppsala, Sweden; Dept of Surgical Sciences, Division of Anesthesiology and Intensive Care, Uppsala University, Uppsala, Sweden; Dept of Medical Sciences, Division of Infectious Diseases, Uppsala University, Uppsala, Sweden; Dept of Medical Sciences, Division of Clinical Chemistry, Uppsala University, Uppsala, Sweden

## Abstract

While fluid resuscitation is fundamental in the treatment of sepsis-induced tissue hypo-perfusion, a sustained positive fluid balance is associated with excess mortality. Crystalloids are the mainstay of fluid resuscitation and use of either synthetic colloids or albumin is controversial. Hyaluronan, an endogenous glycosaminoglycan with high affinity to water, has not been tested as adjuvant in fluid resuscitation.

We sought to evaluate the effects of hyaluronan as an adjuvant to fluid resuscitation in peritonitis induced sepsis. In a prospective, parallel-grouped, blinded model of porcine peritonitis-sepsis, we randomized animals to intervention with adjuvant hyaluronan (add-on to standard therapy) (n=8) or 0.9% saline (n=8). After the onset of hemodynamic instability the animals received an initial bolus of 0.1 % hyaluronan 1 mg/kg/10 min or placebo (saline) followed by a continuous infusion of 0.1% hyaluronan (1 mg/kg/h) or saline during the experiment. We hypothesized that the administration of hyaluronan would reduce the volume of fluid administered (aiming at stroke volume variation <13%) and/or attenuate the inflammatory reaction.

Total volumes of intravenous fluids infused were 17.5 ± 11 ml/kg/h vs. 19.0 ± 7 ml/kg/h in intervention and control groups, respectively (*p* = 0.442). Plasma IL-6 increased to 2450 (1420 – 6890) pg/ml and 3700 (1410 – 11960) pg/ml (18 hours of resuscitation) in the intervention and control groups (NS). In a post-hoc analysis, modified shock index remained lower in intervention group (*p* = 0.011 - 0.037).

In conclusion adjuvant hyaluronan did not reduce the volume needed for fluid administration or decrease the inflammatory reaction. Adjuvant hyaluronan was, however, associated with lower modified shock index. Bearing in mind that the experiment has a limited group-size we suggest that further studies on hyaluronan in sepsis are warranted.

## Introduction

Sepsis is associated with cardiovascular compromise due to absolute and relative hypovolemia, vasodilation, myocardial depression [1] and derangements of the microcirculation [2–4]. While effective fluid resuscitation is essential to antagonize sepsis-induced tissue hypo-perfusion [5], the optimal approach to fluid therapy has not yet been established [6].

A sustained positive fluid balance is associated with higher mortality in septic patients [7–11]. Crystalloids are the first-line fluids recommended for resuscitation in patients with sepsis and septic shock, whereas albumin is recommended in addition when considerable quantities of crystalloids are needed [5]. Although colloids are considered to be more effective volume expanders than crystalloids [12], other colloids than albumin are not recommended for volume resuscitation in sepsis/septic shock [13, 6].

Resuscitation with albumin may be associated with lower total volume administered as compared to crystalloids [14].However, its role as a resuscitation fluid in critical illness [15] or more specifically in sepsis, is not clear [14, 16–19].Albumin is costly, and although it is considered to be safe (as to transmission of pathogens) [20], it is a human blood product with potential side-effects and risks.

Hyaluronan (HA) is a polyanionic, linear glycosaminoglycan, composed of alternating β-d-glucuronate and N-acetyl-β-d-glucosamine [21–23].Due to its large molecular size and negative charges, HA has pronounced hydrophilic and colloid osmotic properties [24–26].The dominating forms of HA *in vivo* have molecular weights of >1000 kDa and are referred to as high molecular weight HA[27] (HMW-HA). HMW-HA is an important constituent of the endothelial glycocalyx layer [27] and is paramount in the maintenance of vascular integrity[28–29].

Inflammation leads to shedding of HA from the vascular endothelial layer [30]. The degradation of HMW-HA is mediated by hyaluronidases [31], as well as by reactive oxygen [32] and nitrogen species [33]. While intact HMW-HA exhibits anti-inflammatory properties [34], degraded, or low molecular weight HA (LMW-HA), has a pro-inflammatory effect [35–36]. HA in plasma has a high turnover rate with a half-life of two to five minutes and is removed primarily by the liver [21]. Elevated levels of plasma HA correlates with more severe disease in critical illness, but studies have rendered conflicting results regarding a possible association with mortality [37–40].

Pretreatment with systemically administered HMW-HA in a rat model of sepsis and mechanical ventilation did not improve macrocirculation, but reduced the inflammatory response in the lung as well as the degree of lung injury [41]. Intravenous administration of HA in humans did not result in any serious adverse events [22]. To the best of our knowledge, systemically administered HMW-HA has not been studied in a context of peritonitis induced sepsis in larger animals or in humans.

We developed, to the best of our understanding, a clinically relevant intensive care, fecal peritonitis/sepsis model in order to test the hypothesis that the administration of HMW-HA would reduce the volume of fluid administered during resuscitation and/or attenuate the hyper-inflammatory state associated with peritonitis induced sepsis.

## Materials and methods

### Animals and ethic statements

The study (protocol: dx.doi.org/10.17504/protocols.io.bwt5peq6) was approved by the Animal Ethics Committee in Uppsala, Sweden (decision 5.8.18-01054/2017, DOUU 2019-014). The care of the animals was carried out in strict accordance with the National Institute of Health guide for the care and use of Laboratory animals (NIH publications No 8023, revised 1978) and all efforts were made to minimize suffering. After premedication and induction of anesthesia, all the animals received continuous intravenous analgesia and were under deep anesthesia. No animal was awake during any moment of the experiment. The study was performed at the Hedenstierna Laboratory, Uppsala University, Sweden.

### Anesthesia and instrumentation

Sixteen pigs (*Sus scrofa domesticus*) of mixed Swedish, Hampshire and Yorkshire breeds of both sexes (mean weight 29.4 ± 1.4 kg) were premedicated with Zoletil Forte^®^ (tiletamine and zolazepam) 6 mg/kg and Rompun^®^ (xylazine) 2.2 mg/kg i.m. After adequate sedation was established we placed the animals in a supine position and introduced a peripheral intravenous catheter in an ear vein. Following a bolus of fentanyl of 5-10 µg/kg i.v., anesthesia was maintained with ketamine 30 mg/kg/h, midazolam 0.1-0.4 mg/kg/h and fentanyl 4 µg/kg/h, in glucose 2.5% during the whole experiment. After adequate depth of anesthesia was assured by absence of reaction to pain stimulus between the front hooves, rocuronium 2.5 mg/kg/h was added as muscle relaxant. Ringer’s acetate was infused i.v. at a rate of 30 ml/kg/h during the first hour and thereafter tapered down to 10 ml/kg/h until the induction of peritonitis.

The animals were under deep anesthesia during the whole experiment (up to 18 hours of sepsis after onset of circulatory instability), including euthanasia. Bolus doses of 100 mg ketamine i.v. were administered if signs of distress or reaction to pain stimulus were noted. In case an animal presented with refractory shock it was euthanized just prior to circulatory collapse (rapidly decreasing systemic arterial pressure, bradycardia and a decrease in end tidal CO_2_).

The animals were tracheostomized and a tube with an internal diameter of eight mm (Mallinckrodt Medical, Athlone, Ireland) was inserted in the trachea and connected to a ventilator (Servo I, Maquet, Solna, Sweden). Thereafter, volume controlled ventilation was maintained as follows: tidal volume (V_T_) 8 ml/kg, respiratory rate (RR) 25/min, inspiratory/expiratory time (I:E) 1:2, inspired oxygen concentration (F_I_O_2_) 0.3 and positive end-expiratory pressure (PEEP) 8 cmH_2_O. The settings of V_T_, I:E and PEEP were maintained constant throughout the protocol. Respiratory rate was adjusted aiming at a PaCO_2_ <6.5 kPa, while F_I_O_2_ was adjusted to keep PaO_2_ >10 kPa.

A triple lumen central venous catheter for fluid infusions and a pulmonary artery catheter (Edwards Life-Science, Irvine CA, USA) for measurement of pulmonary artery pressures and cardiac output (CO) were inserted via the right jugular vein. An arterial catheter for blood pressure measurement and blood sampling was inserted via the right carotid artery. A PiCCO (pulse contour cardiac output) catheter (Pulsion, Munich, Germany) was inserted via the right femoral artery for estimation of stroke volume variation (SVV). Blood gas analyses were performed immediately after sampling and executed on an ABL 3 analyzer (Radiometer, Copenhagen, Denmark). Hemoglobin (hgb) and hemoglobin oxygen saturation were analyzed with a hemoximeter OSM 3 (Radiometer, Copenhagen, Denmark) calibrated for porcine hemoglobin.

We performed a midline laparotomy and catheterized the bladder for urinary drainage. Transit-time flow probe (3 mm, Transonic Systems, Ithaca, New York, USA) was applied around the renal artery. The flow probe was connected to a dual channel flow-meter (T 402, Transonic System Inc, New York, USA) and renal blood flow was recorded continuously. After identification of the caecum a small incision was made, feces was collected and the incision closed. After insertion of a large-bore intra-peritoneal drain, the abdominal incision was closed.

### Study protocol and protocolized resuscitation

#### Preparation of HMW-HA solution

Five grams of HMW-HA 1560 kDa (Sodium hyaluronate Lot# 027362 HA15M-5, Lifecore Biomedical LCC, Chaska, MN, USA) was dissolved in 500 ml 0.9% saline to yield a stock concentration of 1% (10 mg/ml). The solution of 1% HMW-HA 1560 kDa was produced under sterile condition in laminar air-flow, and stored as 50 ml aliquots at −20°C prior to use. On the day of experiment aliquots were thawed and the stock solution was diluted 1:10 in 0.9% saline, to yield 0.1% concentration.

#### Pilot study - kinetics and safety profile of HMW-HA injection

A pilot study was performed prior to the experimental peritonitis model to study simplified plasma kinetics and safety of intravenous HA injection. Animals were injected with HA (HA group, n=3) or 0.9% saline (control group, n=2). Each individual animal in the HA group received a total of six consecutive injections with the 0.1% HMW-HA solution aiming to achieve plasma concentrations of 1000, 5000, 10000, 30000, 50000 and 100000 ng/ml by injecting 0.065, 0.325, 0.65, 1.95, 3.25 and 6.5 ml/kg respectively. The control group received six injections with equal amounts of 0.9% saline. Blood samples (EDTA) for HA analyses were taken at 3 and 45 minutes after each injection and vital parameters were recorded at T=0 (baseline), 5, 10, 15, 25, 35 and 45 minutes.

The experimental time line for the main series is presented in Fig. 1. Baseline measurements were performed after a period of at least 30 min of stabilization following preparation. Peritonitis was established with a peritoneal instillation of autologous feces (2 g/kg body weight in 200 ml warmed 5% glucose solution), after which the large-bore intraperitoneal drain was removed and the abdominal wall closed. The infusion of Ringer’s Acetate was discontinued at the time of the induction of fecal peritonitis.

**Fig 1.**
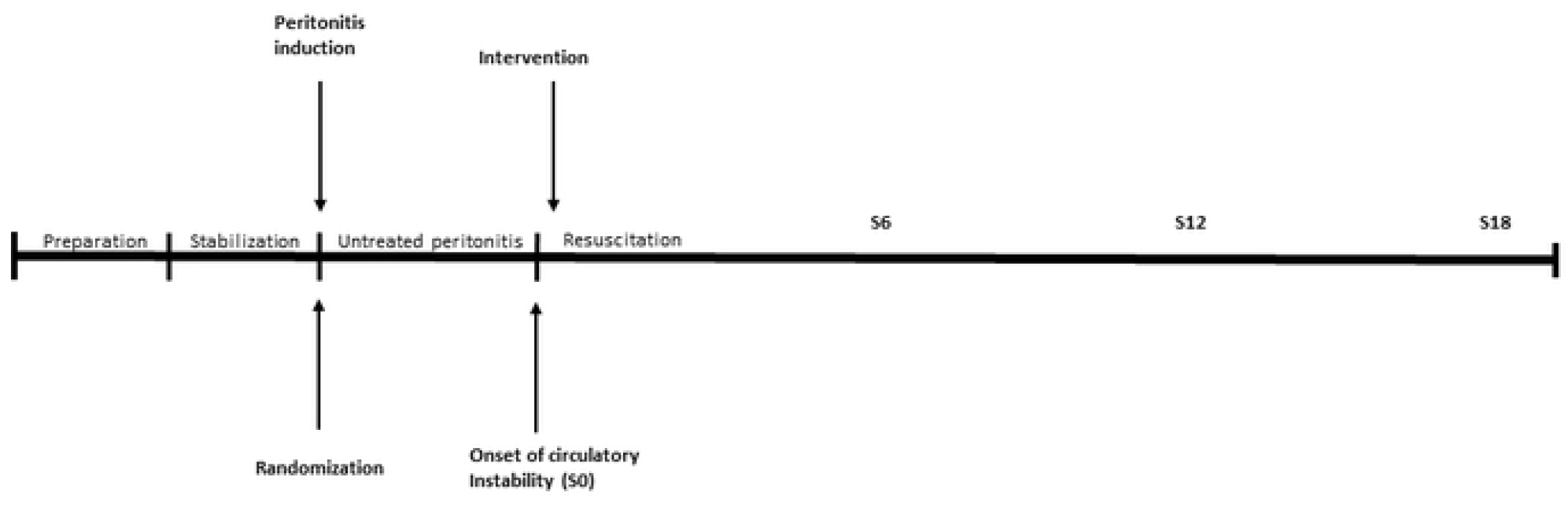
Experimental Time Line. Preparation was followed by at least 30 minutes of stabilization. Thereafter peritonitis was induced. Intervention and resuscitation was initiated at the onset of circulatory instability (S0). S6, S12 and S18 refer to consecutive time points, respectively (hours after S0).

In order to simulate an intensive care setting, the respiratory, circulatory and metabolic maintenance treatments followed a predefined protocol to support vital parameters according to typical invasive monitoring and repeated measurements and sampling. Both intervention and control groups were subject to a protocolized resuscitation that was intitiated at onset of circulatory instability (S0) with Ringer’s Acetate 10 ml/kg/h. The resuscitation protocol was equal to both groups and aimed at MAP > 60 mmHg, guiding fluid and norepinephrine administration by changes in SVV and MAP respectively. If MAP < 60 mmHg and SVV > 15 % fluid was administered, in case of hypotension without increased SVV, infusion of norepinephrine 5 ml/h (40 μg/ml) was started following a bolus of 1 ml (40 μg/ml), and increased stepwise. The fluid therapy consisted of boluses of Ringer’s Acetate of 150 ml, repeated until SVV was steady < 15 %. If MAP was stable > 60 mmHg, infusion was first tapered down to 5 ml/kg/h, and if the animal continued to be stable and SVV maintained < 13% the infusion was stopped [42].

### Experimental design

This was a prospective, parallel-grouped, blinded study with animals randomized (block randomization, sealed opaque envelope) after peritonitis induction into two treatment groups: intervention with HMW-HA (n=8) or control group (n=8). The researchers were blinded for the group allocation until a master file for the whole experiment was produced. After the onset of hemodynamic instability (MAP <60 mmHg for >five min) the intervention group received an initial bolus of 0.1 % HMW-HA solution of 1 mg/kg over ten minutes. The initial bolus was followed by a continuous infusion of the same concentration of 1 mg/kg/h during the rest of the experiment, aiming at a plasma hyaluronan concentration at 10000 - 15000 ng/ml. The control group received the same volume of vehicle (0.9% saline) both as initial bolus and continuous infusion. Immediately after the intervention was initiated, a protocolized resuscitation was started together with Piperacillin/Tazobactam 2 gram in 10 ml of 0.9% saline, given i.v. every 6 hours.

### Analyses and physiologic parameters

We analyzed arterial blood gases at baseline, at the onset of circulatory instability and every hour for the following eighteen hours duration of the experiment. At the same time points, hemodynamic parameters (systemic arterial and pulmonary arterial pressures, CO, heart rate), respiratory parameters (F_I_O_2_, SaO_2_, ETCO_2_, static peak pressure, dynamic and static compliance) and urine output were measured. We calculated modified shock index based on MAP [43,44] and also combined the calculation with hgb and norepinephrine dose. Every three hours mixed venous blood gas analyses was performed, while plasma and urinary samples were collected for analysis every six hours. Stroke volume variation (SVV) was monitored continuously in order to guide fluid resuscitation.

### Cytokine and HA analyses

Porcine-specific sandwich ELISAs were used for the determination of TNF-a, interleukin-6 (IL-6), interleukin-8 (IL-8) and interleukin-10 (IL-10) in plasma (DY690B (TNF-a), DY686 (IL-6), DY535 (IL-8) and DY693B (IL-10), R&D Systems, Minneapolis, MN, USA). The ELISAs had total coefficient of variations (CV) of approximately 6%. Hyaluronan concentration was measured with a commercial ELISA kit (Hyaluronan DuoSet, DY3614, R&D Systems, Minneapolis, MN, USA).

### Tissue analyses post mortem

At the end of the experiment the animals were euthanized with 100 mmol KCl i.v. under deep anesthesia, the skin of the animals was washed with soap, dried with paper and sprayed with ethanol, thereafter the chest wall and abdomen were opened. Tissue samples were collected from left lung (dorsal/basal), heart (left ventricle), liver, spleen, kidney and small intestine. The samples were immersed in 10% buffered formalin immediately. A veterinary pathologist who was blinded for the group allocation evaluated the samples histologically, and inflammatory lesions were graded in a semi-quantitative way: 4, very severe (numerous leukocytes in most parts of the section), 3, severe: (numerous leukocytes in many parts of the section), 2, moderate (moderate numbers of leukocytes diffusely or focally distributed), 1, mild (low number of leukocytes diffusely or focally distributed or 0, lesions were not observed.

Wet-to-dry ratio was measured in samples from the above mentioned locations. Samples were weighed, and dried in an oven, at 50° C, until the weight did not differ between two consecutive measurements.

### Bacterial investigations

Every third hour 0.5 mL arterial blood was collected from a sterile arterial catheter for quantitative blood cultures. Therefrom a 100 µl was cultured on three separate cysteine lactose electrolyte deficient (CLED) agar plates, then cultured at 37°C overnight and colony forming units (CFU) quantified with viable count technique the following day. CFU on only one CLED plate from a time point was interpreted as a contamination otherwise the median of counted CFU/mL was calculated. More than 1 CFU/mL were considered a positive blood culture. Colonies were sent to specification to a MALDI Biotyper (tof-user@FLEX-PC). Samples from lung, spleen and liver were also collected for tissue culture after spraying the organ (surface) with 99% ethanol. App. 1 gram of each tissue was placed in a sterile mortar and mashed in 3 mL saline 0.9%, from where 200 µl was cultured on CLED plates and quantified as described above.

### Statistical analysis

To determine sample size we used data from a previous peritonitis protocol where the fluid balance of the control group had a standard deviation of ± 4 ml/kg/h. Aiming at detecting a difference of 6 ml/kg/h between groups in fluid balance, a power of 0.8 and a significance level of < 0.05 yielded a sample size of eight animals in each group. We tested data for normality by applying the Shapiro-Wilk’s test.

To describe each group separately from baseline to onset of circulatory instability (S0) we used the Student’s t-test, whereas the one-way ANOVA was used to describe the groups separately throughout the experiment. The two-tailed Student’s t-test, the Mann-Whitney U test and the two-way ANOVA were used to compare the two groups, pending distribution of data. Multiple imputation was used in order to replace missing data due to early deaths.

The data are expressed as mean ± SD or median (IQR) as appropriate. We conducted the statistical analyses using SPSS v. 27.0.0 software (SPSS, Inc., Chicago, IL, USA). A *p*-value of < 0.05 was considered to be statistically significant. Bonferroni correction was not used.

The results are presented as n = 8 per group at the baseline, at the onset of circulatory instability (S0), at six (S6) and twelve (S12) hours after onset of circulatory instability, as well as at the end (last observation, prior to imminent death or at 18 hours (S18)). Hourly recordings of hemodynamic and respiratory parameters, as well as blood gas analyses are presented in the electronic supplement (Additional files 1-16). Comparison between the groups over time are presented herein (two-way ANOVA) after performing multiple imputation (i.e. 5) of which p-values are reported as an interval.

## Results

### Pilot study - kinetics and safety profile of HA

The actual measured increase in plasma hyaluronan concentrations followed every injection was 73% (SD ± 13.5%) of the aimed concentration (Supplemental file 17). Hyaluronan removal rate was concentration dependent until the last injection (Supplemental file 18). Plasma hyaluronan concentrations did not return to baseline levels within 45 minutes, resulting in an accumulation effect on the total hyaluronan concentration. The pharmacokinetics of plasma hyaluronan followed a non-linear pattern (Table 1). No adverse effects were observed for either circulatory and respiratory parameters or blood gas analysis. No changes were found for HR, MAP, CO or systemic vascular resistance (SVRI) at any hyaluronan concentration between the hyaluronan and control group (Supplemental file 20). Sub-analysis of HR, MAP, CO over time did not show any consistent changes for any hyaluronan concentration (Supplemental file 21 a-d and 22 a - d).

**Table 1.** Aim vs measured hyaluronan concentration increase after injection.

| Injection hyaluronan (mg/kg) | Aim [hyaluronan] (ng/ml) | Δ hyaluronan increase (ng/ml) |
| --- | --- | --- |
| 0.065 | 1000 | 570 ± 135 |
| 0.325 | 5000 | 3219 ± 3102 |
| 0.65 | 10000 | 7175 ± 2412 |
| 1.95 | 30000 | 21367 ± 2782 |
| 3.25 | 50000 | 48067 ± 18845 |
| 6.5 | 100000 | 79151 ± 8836 |

Netto increase of hyaluronan concentration after each injection with hyaluronan. Values as mean ± SD.

In the main series fourteen out of the sixteen animals survived the experiment until euthanasia (18 hours after onset of circulatory instability), while one animal died of refractory shock during the 18-hours observation period in both treatment (T = S8) and control groups (T = S14).

### Hyaluronan concentration

Plasma hyaluronan concentrations were comparable in the two groups at baseline and at onset of circulatory instability. Plasma hyaluronan concentration (median) increased to 12275, 9060 and 9760 ng/ml during infusion at 6, 12 and 18 hours of the experiment with broad variation. Peritonitis/sepsis *per se* was associated with 3-fold increase of hyaluronan at 18 hours in the control group (Fig 2).

**Figure 2.**
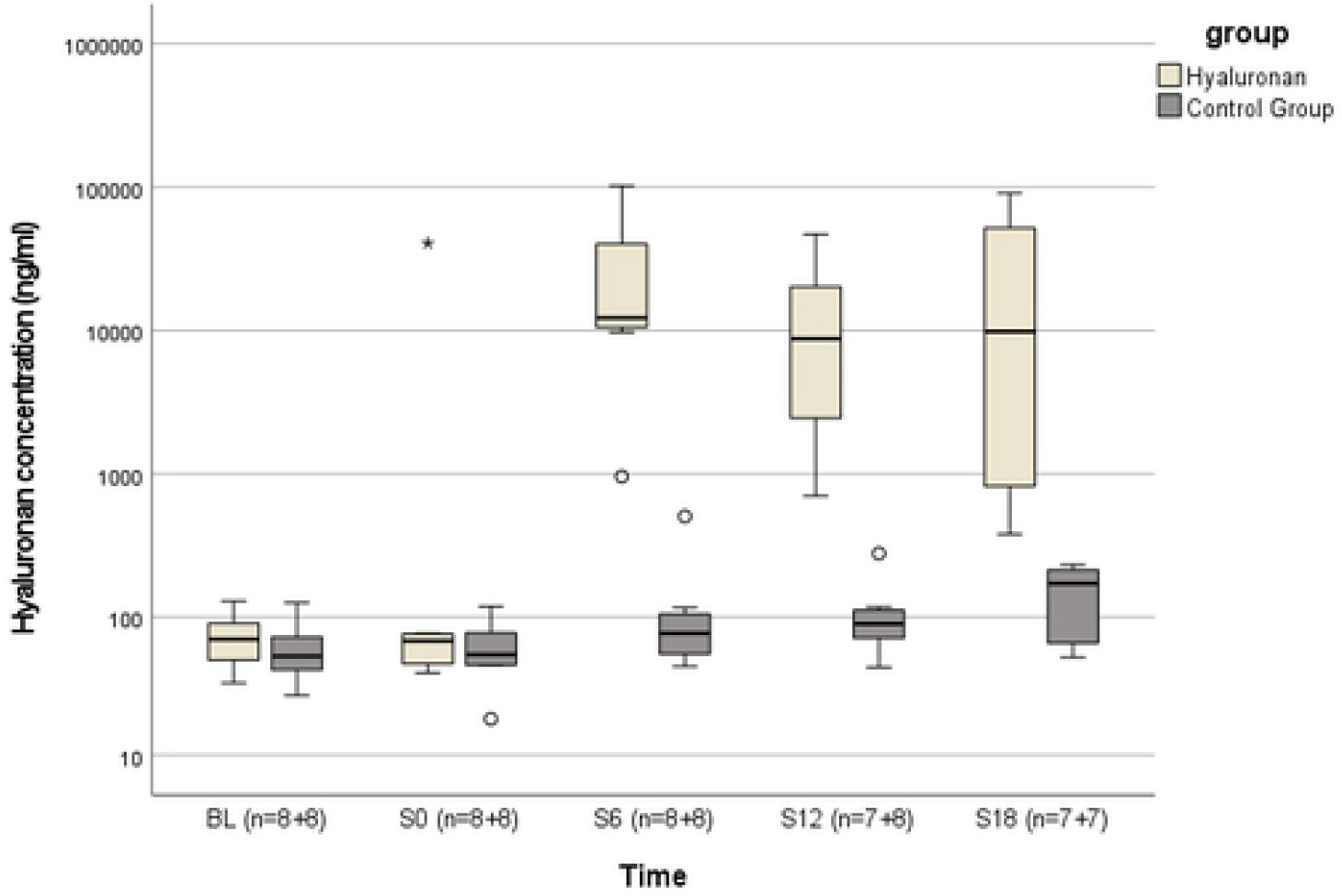
Hyaluronan concentrations. Hyaluronan concentrations in intervention and control groups at baseline, onset of circulatory instability (S0), six (S6), twelve (S12) and eighteen hours (S18) after onset of circulatory instability.

### Hemodynamics

The intervention group presented with circulatory instability (defined as MAP < 60 mmHg > five minutes) within 3.8 ± 1.3 h and the control group within 3.8 ± 1.6 h (*p* = 0.966) from the induction of peritonitis.

The onset of circulatory instability was accompanied by an increase in HR, MPAP, SVV, hgb and temperature in both intervention and control groups (Tables 2 and 3).

**Table 2.**
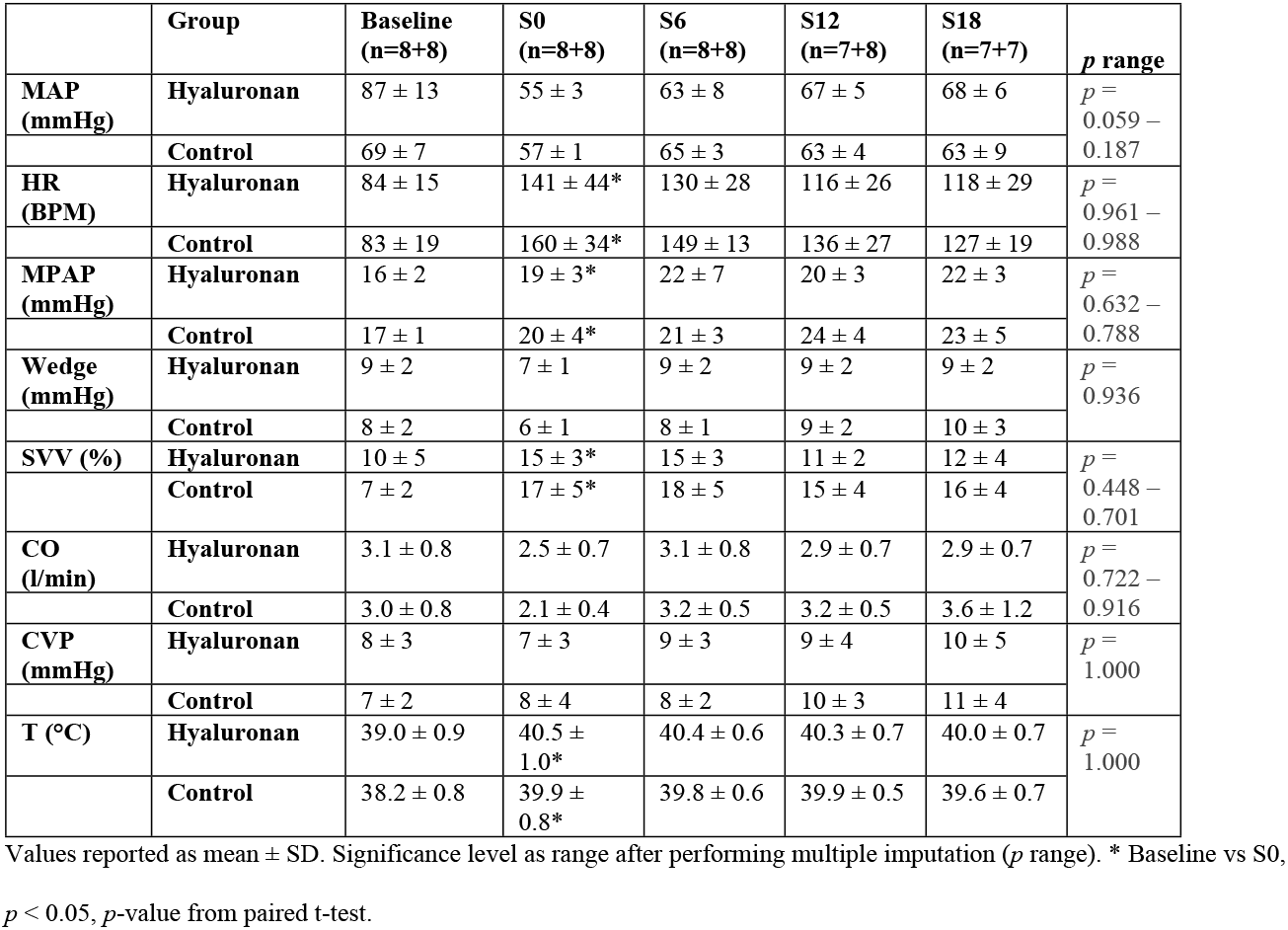
Hemodynamic parameters.

**Table 3.** Arterial blood gas analysis.

|  | Group | BL<br>(n = 8+8) | S0<br>(n = 8+8) | S6<br>(n = 8+8) | S12<br>(n = 7+8) | S18<br>(n = 7+7) | p range |
| --- | --- | --- | --- | --- | --- | --- | --- |
| pH | Hyaluronan | 7.45 ± 0.05 | 7.36 ± 0.07 | 7.38 ± 0.07 | 7.38 ± 0.08 | 7.36 ± 0.12 | $p = 0.931$<br>– 0.990 |
|  | Control | 7.44 ± 0.02 | 7.35 ± 0.04 | 7.37 ± 0.05 | 7.35 ± 0.10 | 7.26 ± 0.17 |  |
| Hgb (g/l) | Hyaluronan | 96 ± 7 | 135 ± 14* | 114 ± 9 | 112 ± 6 | 111 ± 6 | $p = 0.389$<br>– 0.527 |
|  | Control | 93 ± 6 | 139 ± 12* | 122 ± 6 | 113 ± 7 | 107 ± 4 |  |
| Lactate (mmol/l) | Hyaluronan | 2.6 ± 0.8 | 3.4 ± 1.1* | 2.2 ± 1.8 | 1.3 ± 0.4 | 1.3 ± 0.6 | $p = 0.254$<br>– 0.500 |
|  | Control | 2.2 ± 0.5 | 2.5 ± 0.6 | 2.0 ± 0.5 | 2.1 ± 1.4 | 2.4 ± 2.8 |  |
| BE (mmol/l) | Hyaluronan | 3.1 ± 3.2 | - 0.2 ± 4.1 | - 0.7 ± 3.2 | - 0.9 ± 4.3 | - 1.5 ± 6.2 | $p = 0.953$<br>– 0.992 |
|  | Control | 3.9 ± 1.7 | - 0.1 ± 2.5 | - 0.4 ± 2.8 | - 2.7 ± 3.9 | - 5.1 ± 8.9 |  |
| HCO<sub>3</sub><sup>-</sup> (mmol/l) | Hyaluronan | 27.3 ± 2.9 | 24.0 ± 3.6 | 23.7 ± 2.9 | 23.7 ± 3.7 | 23.2 ± 5.2 | $p = 0.955$<br>– 0.987 |
|  | Control | 27.9 ± 1.5 | 23.9 ± 2.1 | 23.9 ± 2.5 | 22.2 ± 3.6 | 20.1 ± 7.2 |  |
| Na (mmol/l) | Hyaluronan | 135 ± 2 | 131 ± 3 | 129 ± 3 | 126 ± 2 | 125 ± 3 | $p = 0.985$<br>– 0.999 |
|  | Control | 135 ± 1 | 131 ± 2 | 129 ± 2 | 126 ± 1 | 125 ± 2 |  |
| K (mmol/l) | Hyaluronan | 3.9 ± 0.3 | 4.9 ± 0.8 | 5.5 ± 0.6 | 5.2 ± 0.5 | 5.0 ± 0.8 | $p = 0.878$<br>– 0.980 |
|  | Control | 3.8 ± 0.4 | 5.0 ± 0.7 | 5.5 ± 0.5 | 5.6 ± 0.4 | 5.5 ± 0.9 |  |

Values reported as mean ± SD. Significance level as range after performing multiple imputation ( $p$ range). \* Baseline vs S0, $p < 0.05$ , $p$ -value from paired t-test.

Lactate increased in intervention group during the same period, but not in the control group (Table 3).

All hemodynamic parameters as well as arterial blood lactate changed comparably in the two groups as a function of time over the length of the resuscitation period (Tables 2 and 3). Neither did the groups differ in regard to wedge pressure, nor CVP as a function of time (Tables 2 and 3).

Modified shock index (HR/MAP) (Fig 3a) and shock index with hgb or with hgb and norepinephrine effects (HR* hgb/MAP and HR*hgb*NE/MAP) (Figs 3b–3c) were comparable at baseline and at the onset of circulatory instability in the two groups. Hyaluronan infusion was associated with lower shock indexes as compared to placebo (Figs 3a – 3c).

**Figures 3a-c.**
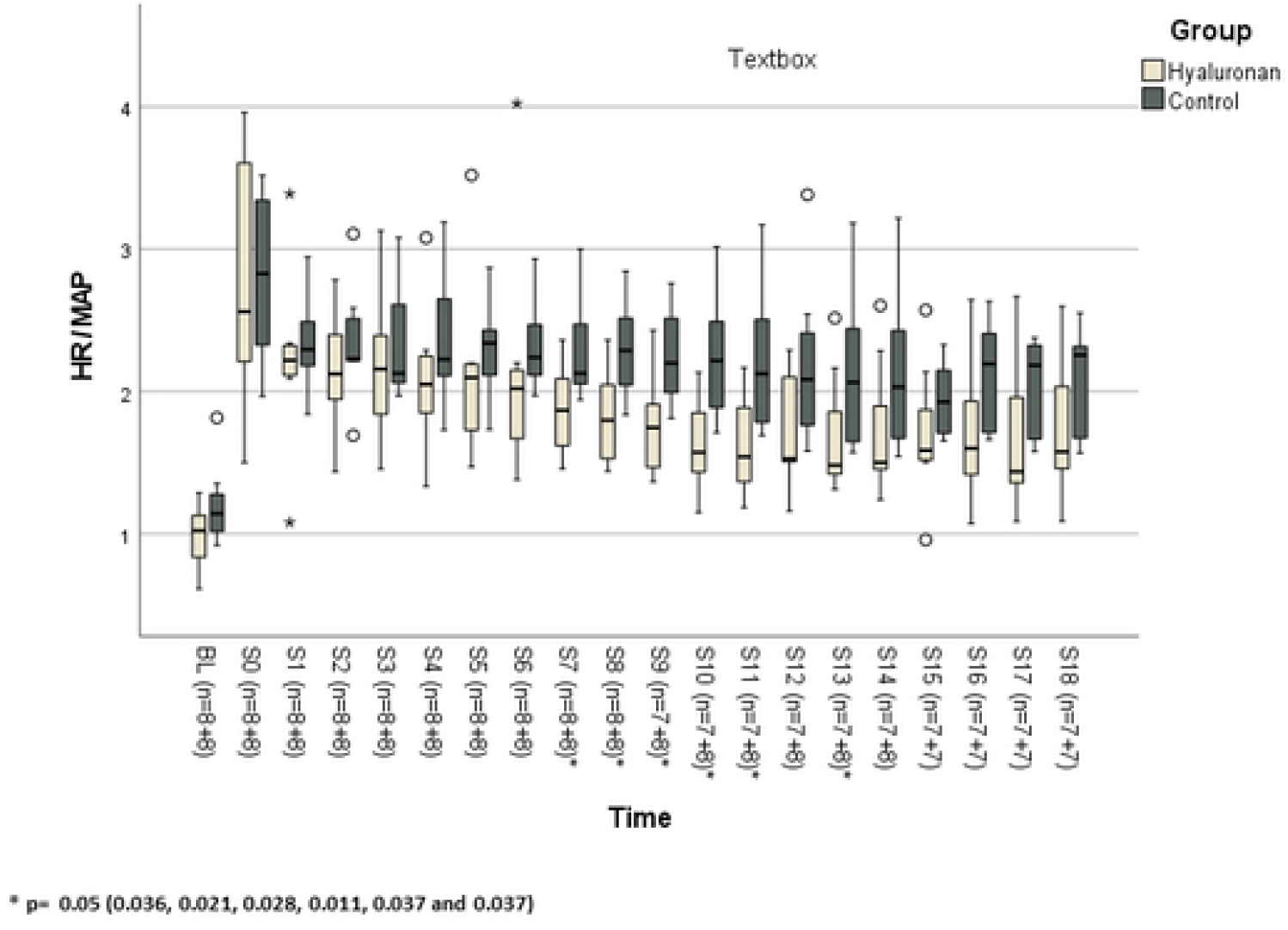
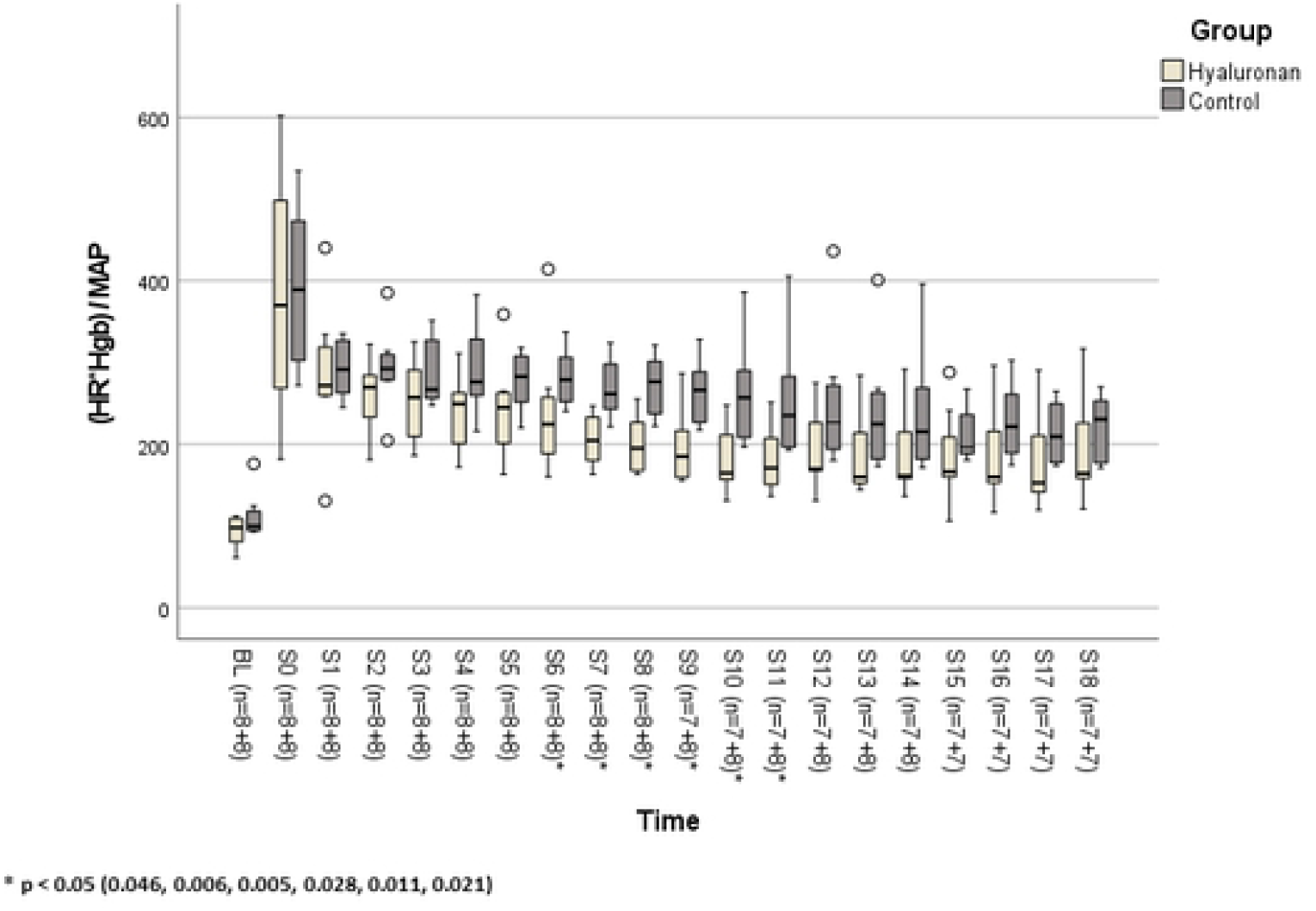
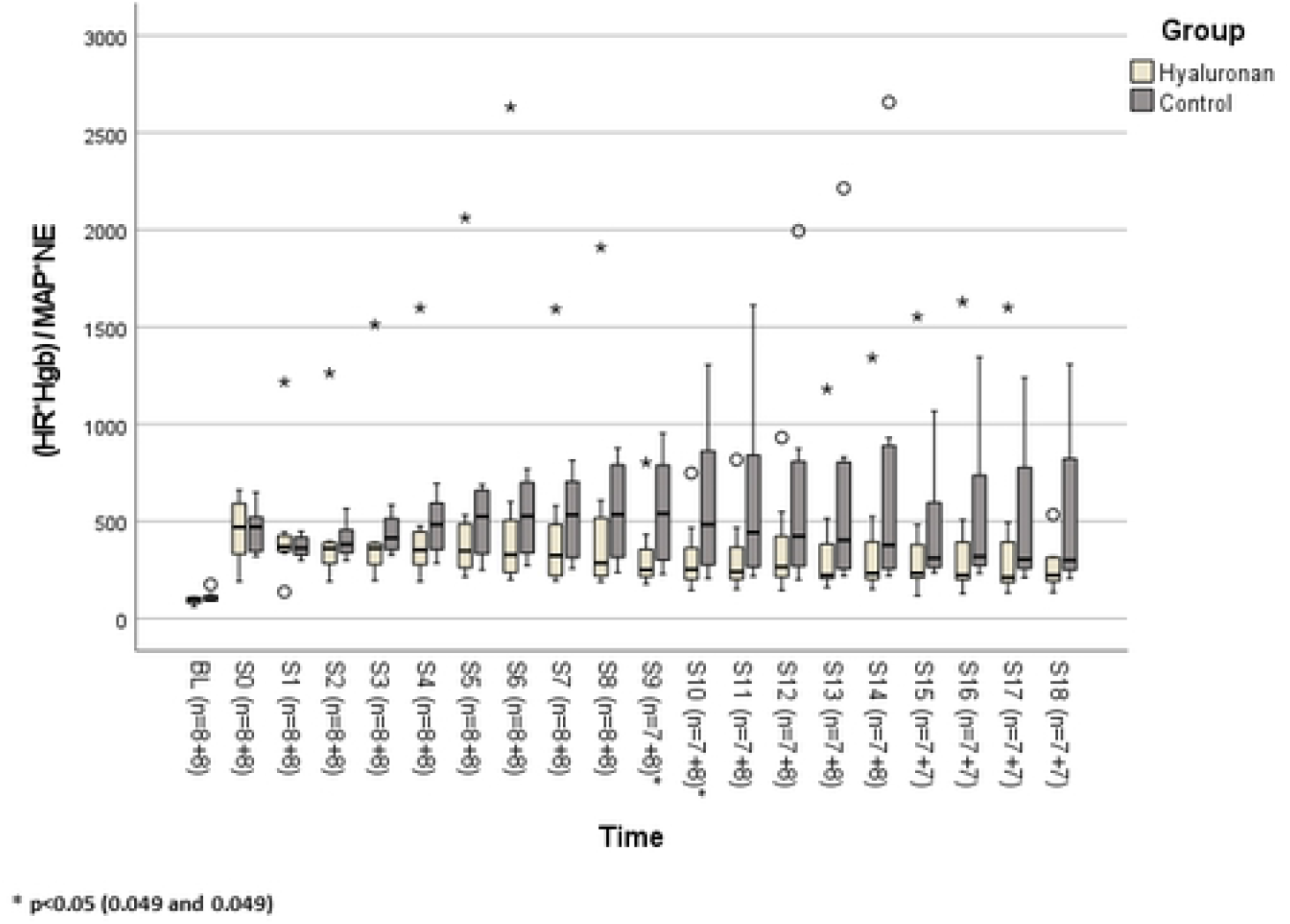
Modified shock indexes. Shock indexes calculated as HR/MAP (3a), HR*hgb/MAP (3b) and HR*hgb*NE/MAP) (3c).

### Respiratory parameters

Onset of circulatory instability was accompanied by comparable decrease of SaO_2_ and P/F ratio from baseline in the two groups. There was a gradual decrease in both dynamic and static compliance in both intervention and control groups respectively, throughout the protocol (Table 4). As RR was adjusted (intervention group: range 25 – 50/min) (control group: range 25 – 40/min) in order to maintain normocapnia we observed that PEEP_TOT_ was stable at 8 cm H_2_O in the both groups throughout the experiment. A progressive increase in static peak pressure throughout the protocol was statistically significant in the control group (*p* = 0.001 – 0.004) but not in the intervention group (*p* = 0.350 – 0.594) throughout the resuscitation period.

**Table 4.**
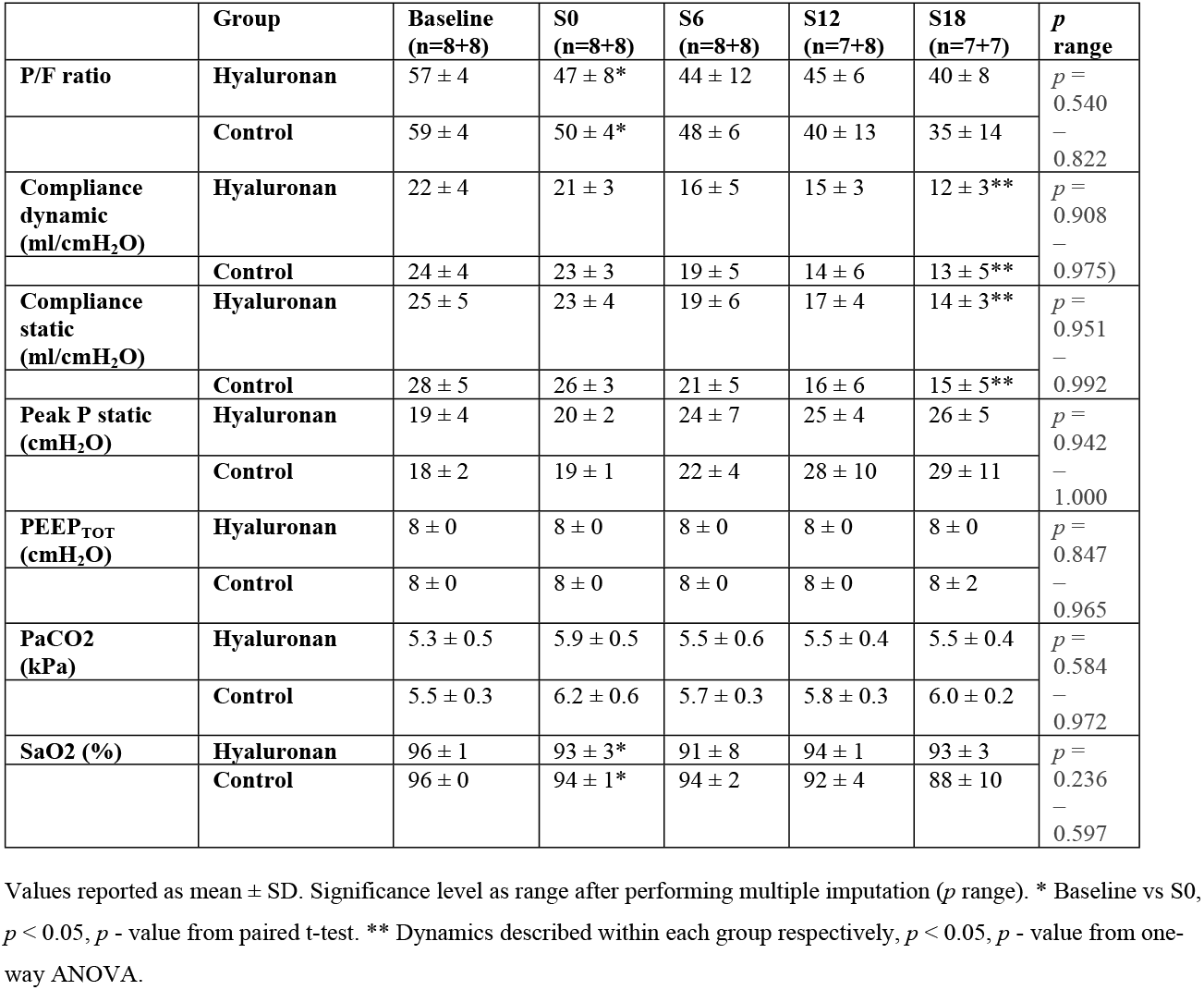
Respiratory parameters.

### Fluid balance, norepinephrine dosage, kidney function and electrolytes

Total volumes of fluid administered during the experiment were 17.5 ± 11 ml/kg/h vs. 19.0 ± 7 ml/kg/h in intervention and control groups, respectively (*p* = 0.442). Weight gain was 12.5 ± 3.1 kg in the intervention and 14.0 ± 2.3 kg in the control group (*p* = 0.328). The average norepinephrine dosage was 1.2 ± 1.6 µg/kg/min and 1.0 ± 0.8 µg/kg/min in the intervention and the control groups, respectively (*p* = 0.721) (Additional file 23).

Urine production decreased from baseline to onset of circulatory instability from 4.1 ± 4 ml/kg/h to 0.5 ± 0.4 ml/kg/h (*p* = 0.045) and from 3.0 ± 2 ml/kg/h to 0.5 ± 0.4 ml/kg/h (*p* = 0.015) in the intervention and control groups, respectively. Hourly diuresis was comparable in the two groups throughout the resuscitation period, in average 2.1 ± 1.3 ml/kg/hour in the intervention and 1.7 ± 0.9 ml/kg/hour in the control group (*p* = 0.442) (Additional file 25).

Renal arterial blood flow decreased from baseline to onset of circulatory instability in both intervention and control group: from 132 ± 79 ml to 61 ± 46 ml (*p* = 0.001) vs. from 138 ± 63 ml to 72 ± 34 ml (*p* = 0.002). Blood flow changed comparably in the two groups as a function of time (*p* = 0.873 – 0.976) throughout the protocol (Additional file 26).

Plasma creatinine increased from baseline 77 ± 17 µmol/l to 100 ± 19 µmol/l at onset of circulatory instability in intervention group and from 72 ± 12 µmol/l to 92 ± 14 µmol/l in control group, with comparably increasing plasma concentrations throughout the resuscitation period in both groups respectively as a function of time. Creatinine clearance is depicted in additional file 27. Plasma urea, whole blood Na, whole blood K, BE and HCO_3_^-^ followed a similar pattern over time in the two groups (Tables 3 and 5).

**Table 5.**
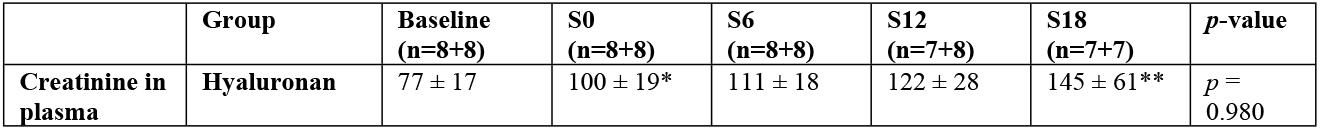
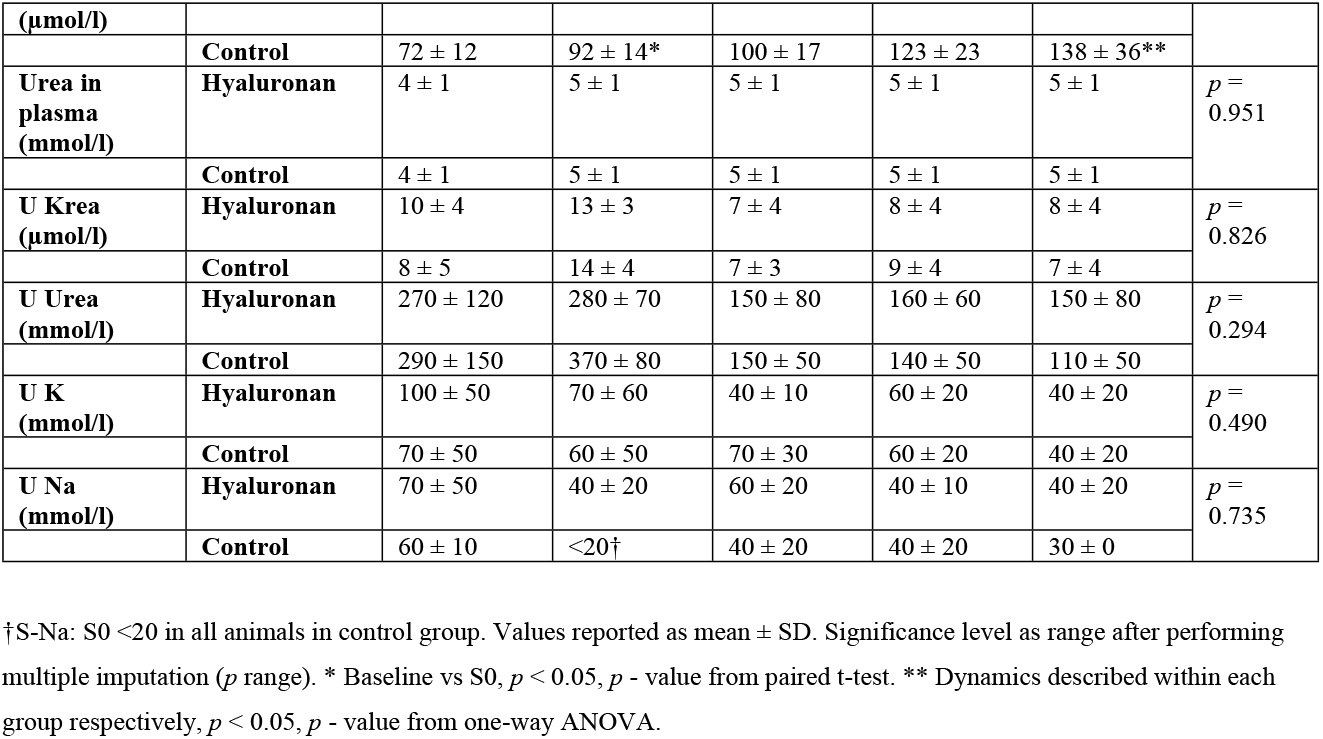
Creatinine and urea in plasma and urine markers.

Finally, there were no differences between groups in urine creatinine, urea, sodium, or potassium concentrations between groups as a function of time (Table 5).

### Cytokines

The concentration of IL-6, IL-8, IL-10 and TNF-α in plasma increased from baseline to onset of circulatory instability (S0) in both intervention and control groups. The dynamics in cytokine concentrations in plasma were comparable in the two groups throughout the experiment (Table 6).

**Table 6.** Cytokines in plasma.

| Cytokine | Group | Baseline (n=8+8) | S0 (n=8+8) | S6 (n=8+8) | S12 (n=8+8) | S18 (n=7+8) | $p$ range |
| --- | --- | --- | --- | --- | --- | --- | --- |
| **IL-6 (pg/ml)** | **Hyaluronan** | 150 (-10 – 470) | 5650 (3070 – 6540)* | 4030 (2670 – 6570) | 4000 (1730 – 8380) | 2450 (1420 – 6890) | $p = 0.614 - 0.859$ |
|  | **Control** | 270 (130 – 460) | 5340 (3460 – 6870)* | 3700 (2050 – 6950) | 3910 (2300 – 8330) | 3700 (1410 – 11960) |  |
| **TNF-α (pg/ml)** | **Hyaluronan** | 150 (100 – 310) | 240 (140 – 420) * | 170 (90 – 320) | 160 (130 – 190) | 160 (140 – 250) | $p = 0.932 - 0.957$ |
|  | **Control** | 170 (90 – 380) | 230 (140 – 390)* | 140 (100 – 250) | 160 (120 – 230) | 150 (110 – 240) |  |
| **IL-10**<br>**(pg/ml)** | **Hyaluronan** | 240 (-20 – 760) | 820 (450 – 1350)* | 860 (690 – 1480) | 660 (430 – 1140) | 500 (360 – 770) | $p = 0.748 - 0.796$ |
|  | **Control** | 330 (220 – 570) | 610 (490 – 820)* | 720 (570 – 1180) | 600 (540 – 910) | 540 (380 – 790) |  |
| **IL-8**<br>**(pg/ml)** | **Hyaluronan** | 60 (50 – 80) | 130 (90 – 150)* | 120 (80 – 190) | 90 (50 – 190) | 80 (40 – 170) | $p = 0.694 - 0.741$ |
|  | **Control** | 60 (50 – 90) | 110 (80 – 150)* | 110 (70 – 130) | 80 (60 – 110) | 60 (40 – 130) |  |

One animal died before finishing the protocol in both groups, last sample before imminent death is included in the analyses of the time point after death, that is S12 for the animal that died in group 1 (204 at S8), and S18 for the animal that died in group 2 (207 at S14), before finishing the protocol. Values as median (95% CI). \* Baseline vs S0, $p < 0.05$ , $p$ - value from paired t-test.

### Wet-to-dry ratio

Wet-to-dry ratios at the end of the experiment were comparable in the two groups (Additional file 28).

### Blood and tissue cultures

#### Blood cultures

All animals, except one in the intervention group, had positive blood cultures at some time point during the experiment (Additional file 29). Number of CFU/ml (*p* = 0.682) were comparable in the two groups throughout the experiment. Positive cultures were of mixed etiology, with a dominance (> 90%) of E. coli.

#### Tissue cultures

All animals had viable bacteria in at least one organ (lung, liver, spleen). All three tissue cultures had viable bacteria in five animals in intervention group and in four animals in the control group. The two groups did not differ in CFU/g in either of the tested organs.

Cultures of lung tissue had viable bacteria in five animals in intervention group (median: 50, IQR 475) and in six animals in the control group (median 2: 200, IQR 4400) (*p* = 0.382). Liver tissue cultures had viable bacteria in six animals in intervention group (median: 254, IQR 405) and in five animals in control group (median: 236, IQR 1682) (*p* = 1). Tissue cultures of spleen had viable bacteria in all animals in intervention group (median: 5174, IQR 15692) and in all but one animal in the control group (median: 1813, IQR 8107) (*p* = 0.195).

### Histology

Lung samples showed acute inflammatory lesions in samples from four animals of the intervention group and seven animals of the control group, the lesions varied in intensity between individual animals. Lung lesions were comparable in intervention (median 1, IQR 2) and control groups (median 2, IQR 2) (*p* = 0.234) (Figs 4a and 4b).

**Figures 4a and 4b.**
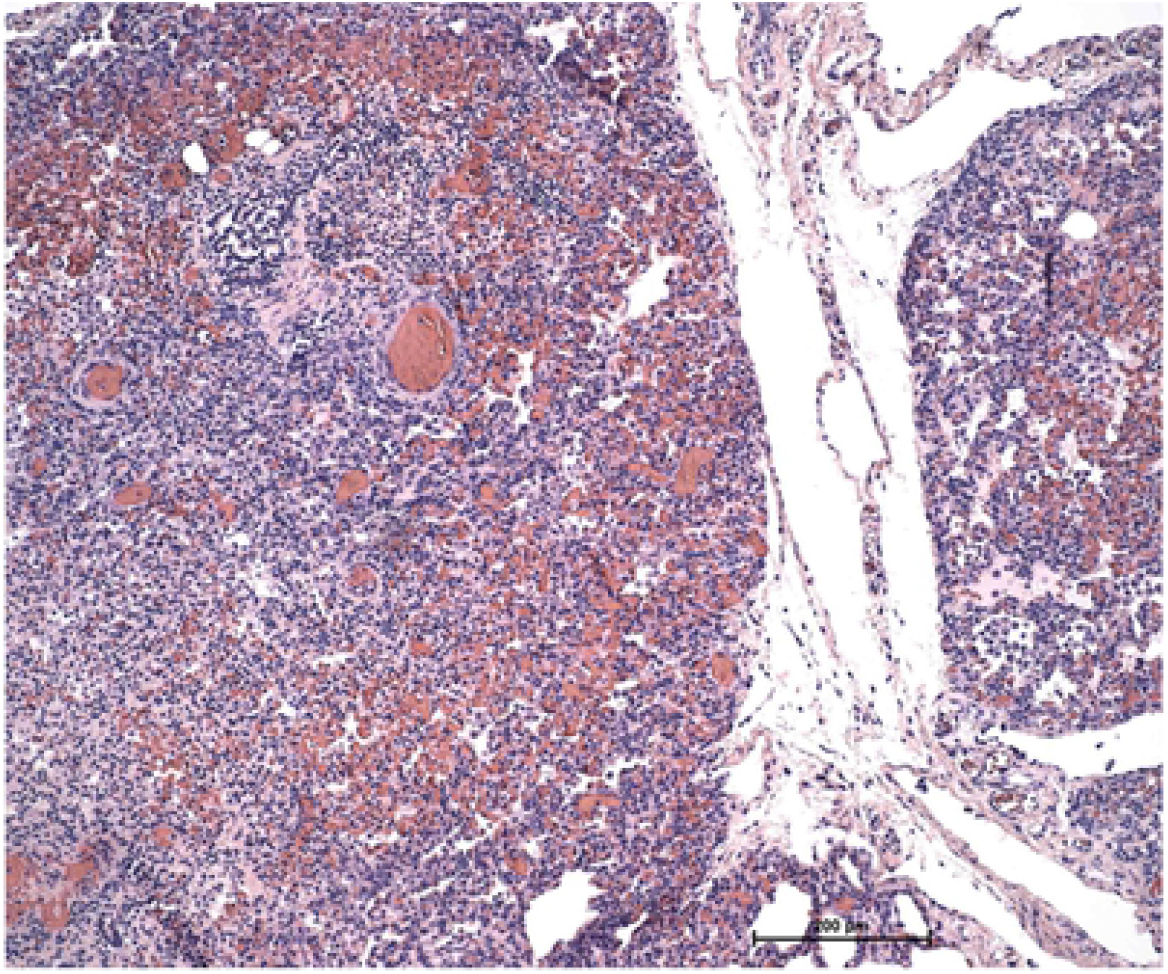
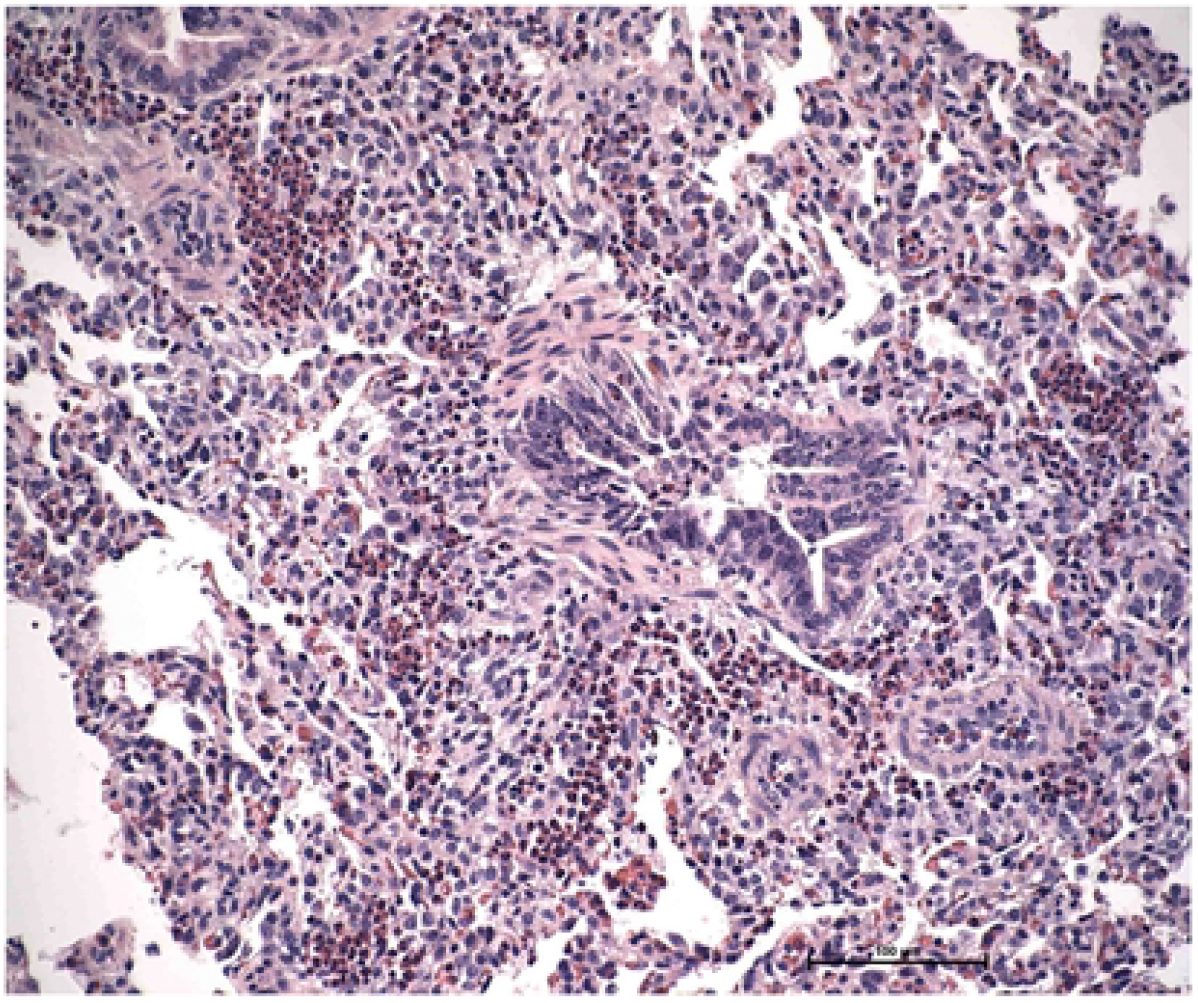
Lung histology. Lung histology, representative samples of lung lesions from one animal in intervention and control groups, respectively (please note the different magnification).

No significant lesions were visualized in heart tissue in either group. All but one animal in each group had acute focal/multifocal degeneration in the liver and coagulative necrosis of hepatocytes. Liver lesions were comparable between groups: intervention group (median 3, IQR 2) and control groups (median 1, IQR 1) (*p* = 0.065). In intestinal samples, the epithelial lining was generally preserved, but the gut mucosa in all pigs was infiltrated with mixed leucocytes to a varying degree. There was no difference between intervention (median 2, IQR 1) and control groups (median 2, IQR 2) (*p* = 0.328) as to inflammatory lesions of the intestine (Figs 5a and 5b).

**Figures 5a and 5b.**
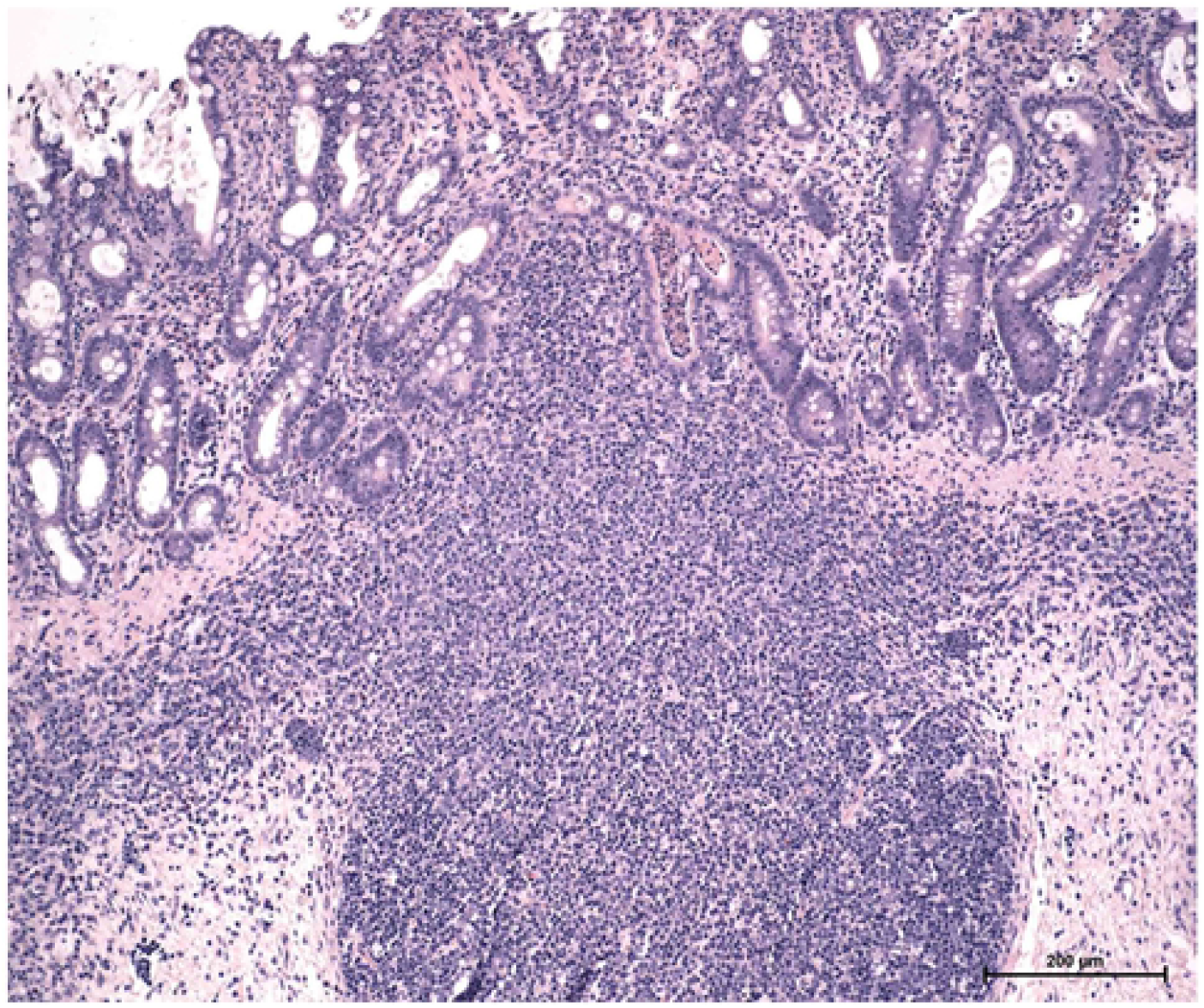
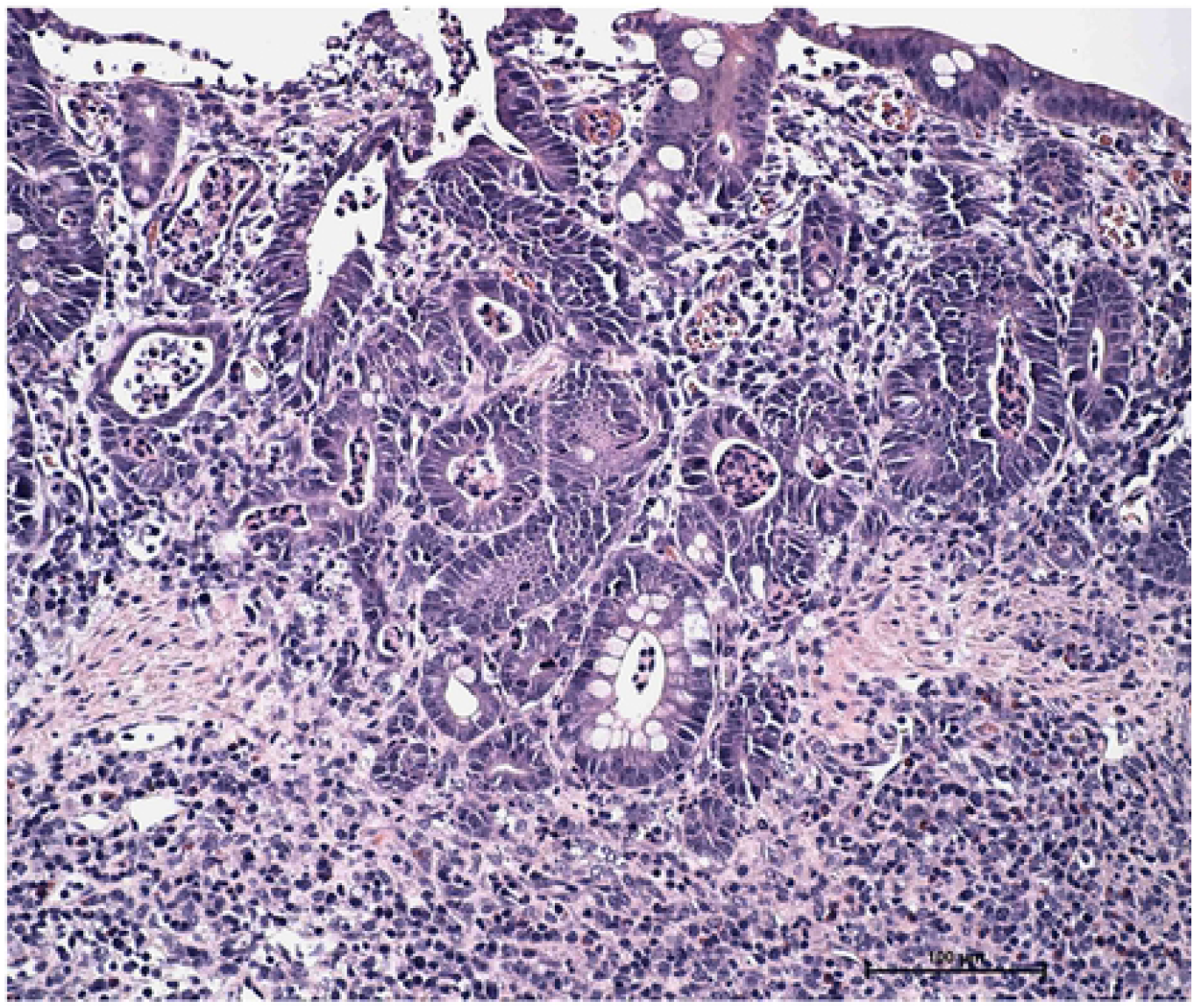
Histology of intestine. Histology of intestine, representative samples of intestinal lesions from one animal in intervention and control groups, respectively (please note the different magnification).

Lesions in kidney samples were rare (intervention: median 0, IQR 1, control group: median 0, IQR 1) and equally distributed between groups (*p* = 0.959). Three animals in the intervention group (median 0, IQR 3) and three animals in the control group (median 0, IQR 3) exhibited inflammatory lesions of the spleen, manifested as acute purulent inflammation of the splenic capsule. The capsular inflammation generally extended to the parenchyma in subcapsular areas. There was no difference between groups (*p* = 1.000) regarding inflammatory lesions of the spleen.

## Discussion

The main finding of the present study was that, contrary to our hypothesis, high molecular weight hyaluronan infusion did not decrease the total volume of fluid resuscitation in the early phase of peritonitis-induced sepsis. Weight gain, urine production, tissue wet-to-dry ratios and histology in the intervention and control groups were comparable. Furthermore, plasma cytokine concentrations were comparable over the length of the experiment. In the post hoc analyses, on the other hand, modified shock index with and without the additional combined effect of hemoglobin suggested that HMW-HA in the present model was associated with less hemodynamic instability.

We used a porcine model of fecal peritonitis to mimic sepsis and septic shock in patients. All the animals presented with circulatory instability after a period of untreated peritonitis and some developed shock/refractory shock, although the disease severity was not as pronounced as previously described by us. The heterogeneous nature of the model is consistent with the sepsis panorama seen in patients, where a similar infectious stimulus results in resolving infection in some individuals while others develop a severe shock state with hyperlactatemia and resistance to resuscitation. Tissue and blood cultures as well as the dynamics in plasma concentrations of the measured cytokines (IL-6, IL-8, IL-10 and TNF-α) were all consistent with a severe infection and the systemic inflammation as seen in sepsis [45].

In the present study isolated hemodynamic parameters (HR, MAP, SVV, CO, mixed venous saturation or lactate), body weight gain (kg before vs. after the experiment), wet-to-dry ratio and fluid balance did not differ between groups, suggesting that systemically administered HMW-HA do not exert any volume sparing effects in sepsis resuscitation. However, previously described shock indexes may predict morbidity and mortality better than isolated hemodynamic parameters [46,47], including the modified shock index[43,44]. In the current study neither HR nor MAP differed between groups, however a post hoc analysis of modified shock index (HR/MAP) revealed a statistically significant difference between groups at several time points with a less pronounced hemodynamic instability in the intervention group, this held true also when including hemoglobin and norepinephrine dosage in the calculation.

The discussion about the optimal fluid for volume resuscitation in sepsis and septic shock continues [6]. While fluid administration is indispensable in the resuscitation phase of sepsis and early fluid administration is associated with a better outcome [5], fluid overload is associated with increased mortality [7–11].Intravenous colloids oppose the transcapillary fluid flux [48] and are more effective volume expanders than crystalloids [12]. However this advantage in sparing volume of total fluid administered during resuscitation is not associated with better outcome in critical illness [14].

HA is a molecule with pronounced hydrophilic properties that plays a significant role in the regulation of water homeostasis and is present in all tissues and body fluids [21]. It is an important contributor to colloid osmotic pressure in, for instance, synovial fluid [24]. Solutions of HA has highly non-ideal characteristics regarding colloid osmotic pressure, implying a rapidly increasing colloid osmotic pressure with increasing concentration [49], with the colloid osmotic pressure of HA exceeding that of albumin at a concentration of 1 mg/ml respectively [50]. The colloid osmotic pressure of solutions containing mixtures of HA and albumin exceed the sum of the colloid osmotic pressures of each solution separately, at equal concentrations [51].

In the kinetics and safety pre-study, HMW-HA was administered intravenously in healthy pigs without any observed negative effects during the experiment. The measured hyaluronan concentration in plasma was in average 73 ± 13.5% of the aimed concentration, approximating blood volume in the pig to be 65 ml/kg. From these data we calculated that a plasma concentration of 15 000 ng/ml would be achieved with an initial bolus of 1 mg/kg followed by an infusion of 1mg/kg/h. Doses as high as 1.5 - 12 mg/kg have been administered without any serious adverse events in healthy humans [22].

Plasma concentrations of HA were comparable in the intervention and control groups at baseline and at onset of circulatory instability. Peritonitis/sepsis was associated with 3-fold increase of hyaluronan in the control group during the resuscitation period. The intervention group showed a greater increase of plasma HA due to the infusion of exogenous hyaluronan, and values were considerably higher in the intervention group than values previously reported in sepsis [38,39,29,40]. Since this was associated with less hemodynamic instability (modified shock indexes) our findings may support the notion that HA exerts a protective effect in critical illness, as suggested previously [39].

Neither the histological analysis, bacterial cultures or the cytokine response in plasma revealed a difference in inflammation between the two groups, suggesting that HA administered in sepsis, after onset of circulatory instability, does not counteract the state of hyper-inflammation associated with the early phase of peritonitis induced sepsis. HMW-HA is anti-inflammatory and immunosuppressive [34]. It reduces the pro-inflammatory cytokine response in plasma [52], promotes resolution of infection [53] and potentially renders antibodies a higher neutralizing capacity through steric exclusion [21]. Fractions of HA, or low molecular weight HA (250 kDa), on the other hand, is a strong signal of tissue damage [54] and induces a pro-inflammatory response through several pathways [35,52,36] with production of pro-inflammatory cytokines and enhanced secretion of nitric oxide [53]. The present study does not support the previous notion that HMW-HA has anti-inflammatory effects.

Our study has several limitations. Most importantly it is an animal model which implies possible species differences in host response, both in regard to infectious insult and to intervention, as compared to humans. It is also a study of limited size and small differences between groups might not have been detected, even more so since the peritonitis model used presents with a heterogeneous panorama of disease severity. Furthermore peritoneal lavage was not performed for source control. Several pigs had positive blood cultures already at baseline, a finding most likely explained by the fact that baseline measurements were performed after laparotomy, “pre-baseline” blood cultures immediately at arrival should be considered for future protocols. There are also open questions about optimal concentration, timing [55] and mode of administration of a HA solution. Administration of rapid crystalloid infusions increases plasma concentration of HA and might disrupt the glycocalyx [56]. Thus, the fact that we started an infusion of HMW-HA at onset of circulatory instability alongside with resuscitation with crystalloid (infusion and boluses as needed) might have interfered with the intervention. Limiting fluid resuscitation in the intervention group to nothing but HMW-HA solutions could be an alternative approach to study the potential benefit of HMW-HA solutions in sepsis resuscitation.

## Conclusions

In conclusion, the current study does not support the hypothesis that HMW-HA reduces the volume of fluid administered during resuscitation and/or attenuate the hyper-inflammatory state associated with peritonitis induced sepsis. However, a post-hoc analysis of modified shock index showed that systemically administered HMW-HA is associated with a less pronounced hemodynamic instability as to controls. This finding, albeit implied with the potential drawbacks of post-hoc analyses *per se*, suggests that a beneficial role of HMW-HA in the context of sepsis resuscitation is possible.

## Supporting information captions

**S1. Mean arterial pressure (MAP)**. Hourly recordings of MAP throughout the experiment, intervention and control group.

**S2. Heart rate (HR) beats per minute (BPM)**. Hourly recordings of HR throughout the experiment, intervention and control group.

**S3. Hemoglobin (hgb) g/l**. Hourly analyses of hgb throughout the experiment, intervention and control group.

**S4. Lactate mmol/l**. Hourly analyses of lactate throughout the experiment, intervention and control group.

**S5. pH**. Hourly analyses of blood pH throughout the experiment, intervention and control group.

**S6. Base excess (BE) mmol/l**. Hourly analyses of BE throughout the experiment, intervention and control group.

**S7. HCO_3_**- **mmol/l**. Hourly analyses of blood HCO_3_- throughout the experiment, intervention and control group.

**S8. Mixed venous SaO**_**2**_ **%**. Analyses of mixed venous SaO_2_ every three hours throughout the experiment, intervention and control group.

**S9. Mean pulmonary arterial pressure (MPAP) mmHg**. Hourly recordings of MPAP throughout the experiment, intervention and control group.

**S10. Stroke volume variation (SVV) %**. Hourly registrations of SVV, continuously recorded throughout the experiment, intervention and control group.

**S11. Cardiac output (CO) l/min**. Hourly recordings of CO throughout the experiment, intervention and control group.

**S12. Central venous pressure (CVP) mmHg**. Hourly recordings of CVP throughout the experiment, intervention and control group.

**S13. Temperature T °C**. Hourly recordings of T throughout the experiment, intervention and control group.

**S14. PaO**_**2**_ **to inspired O**_**2**_ **fraction (P/F-ratio)**. Hourly recordings of P/F-ratio throughout the experiment, intervention and control group.

**S15. Static compliance (Cstat) ml/cmH**_**2**_**O**. Hourly recordings of Cstat throughout the experiment, intervention and control group.

**S16. Static peak pressure cmH**_**2**_**O**. Hourly recordings of static peak pressure throughout the experiment, intervention and control group.

**S17. Plasma concentration of hyaluronan ng/ml – pilot study**. Aimed hyaluronan plasma concentrations vs measured (ng/ml) after injection of 0.065, 0.325, 0.65, 1.95, 3.25 and 6.5 mg/kg respectively.

**S18. Hyaluronan kinetics 1**. Plasma concentration of hyaluronan, aim vs directly after bolus injection as well as after 45 minutes, hyaluronan removal from plasma in ng/ml/min, values as mean ± SD.

**S19a. Hyaluronan kinetics 2**. Estimation of amount hyaluronan required to reach goal concentration of 10000 ng/ml according to blood volume of pigs (65 ml/kg) and goal concentration 15000 ng/ml.

**S19b. Hyaluronan kinetics 3**. Estimation of amount hyaluronan required to reach goal concentration of 10000 ng/ml according to pilot study.

**S20. Hemodynamic changes after injection of hyaluronan or 0.9% saline at several time points**. Changes are compared with baseline values for each individual experiment.

**S21 a-d. Changes in heart rate (HR) BPM, mean arterial pressure (MAP) (mmHg), cardiac output (CO) l/min and systemic vascular resistance index (SVRI) (dynes) compared with baseline**. Heart rate (HR), mean arterial pressure (MAP), cardiac output (CO) and systemic vascular resistance index for different hyaluronan concentrations. Δ indicates the change in HR/MAP/CO/SVRI after injection with hyaluronan compared with the baseline value. No differences were found between the hyaluronan and control group. *Δ HR for HA 50000 is considered as outliner.

**S22 a-d. Changes in heart rate (HR) BPM, mean arterial pressure (MAP) (mmHg), cardiac output (CO) l/min and systemic vascular resistance index (SVRI) (dynes) in % as compared to baseline**. HR, MAP, CO and SVRI for different hyaluronan concentrations. Δ indicates the change in HR/MAP/CO/SVRI after injection with hyaluronan compared with the baseline value in %. No differences were found between the hyaluronan and control group. *Δ HR for HA 50000 is considered as outliner

**S23. Norepinephrine administration µg/kg/min**. Administration of norepinephrine presented hourly throughout the experiment, intervention and control group.

**S24. Fluid administration ml/h**. Registration of fluid administration throughout the experiment, presented hourly, intervention and control group.

**S25. Diuresis ml/kg**. Hourly measurements of diuresis throughout the experiment, intervention and control group.

**S26. Flow arteria renalis % of flow at baseline**. Hourly recordings of flow arteria renalis throughout the experiment, intervention and control group.

**S27. Creatinine clearance ml/h**. Analyses of creatinine clearance every six hours throughout the experiment, intervention and control group.

**S28. Wet-to-dry ratio**.

**S29. Blood cultures**. CFU/ml. Data presented as Median (IQR).

